# Formation of Müller glia-derived progenitor cells in retinas depleted of microglia

**DOI:** 10.1101/2023.06.08.544205

**Authors:** Heithem M. El-Hodiri, James Bentley, Alana Reske, Isabella Palazzo, Warren A. Campbell, Nicklaus R. Halloy, Andy J. Fischer

## Abstract

Recent studies have demonstrated the complex coordination of pro-inflammatory signaling and reactive microglia/macrophage on the formation Müller glial-derived progenitor cells (MGPCs) in the retinas of fish, birds and mice. We generated scRNA-seq libraries to identify transcriptional changes in Müller glia (MG) that result from the depletion of microglia from the chick retina. We found significant changes in different networks of genes in MG in normal and damaged retinas when the microglia are ablated. We identified a failure of MG to upregulate Wnt-ligands, Heparin binding epidermal growth factor (HBEGF), Fibroblast growth factor (FGF), retinoic acid receptors and genes related to Notch-signaling. Inhibition of GSK3β, to simulate Wnt-signaling, failed to rescue the deficit in formation of proliferating MGPCs in damaged retinas missing microglia. By comparison, application of HBEGF or FGF2 completely rescued the formation of proliferating MGPCs in microglia-depleted retinas. Similarly, injection of a small molecule inhibitor to Smad3 or agonist to retinoic acid receptors partially rescued the formation of proliferating MGPCs in microglia-depleted damaged retinas. According to scRNA-seq libraries, patterns of expression of ligands, receptors, signal transducers and/or processing enzymes to cell-signaling via HBEGF, FGF, retinoic acid and TGFβ are rapidly and transiently upregulated by MG after neuronal damage, consistent with important roles for these cell-signaling pathways in regulating the formation of MGPCs. We conclude that quiescent and activated microglia have a significant impact upon the transcriptomic profile of MG. We conclude that signals produced by reactive microglia in damaged retinas stimulate MG to upregulate cell signaling through HBEGF, FGF and retinoic acid, and downregulate signaling through TGFβ/Smad3 to promote the reprogramming on MG into proliferating MGPCs.

## Introduction

The process of retinal regeneration varies greatly between vertebrate species. In fish retinas neuronal regeneration is a robust process that restores visual function following injury, whereas this process is far less robust in birds and absent in mammals (Hitchcock and Raymond, 1992; Karl et al., 2008; Raymond, 1991). Müller glia (MG) have been identified as the cell of origin for progenitors in regenerating retinas (Bernardos et al., 2007; Fausett and Goldman, 2006; Fausett et al., 2008; Fischer and Reh, 2001a; Ooto et al., 2004). In normal retinas MG are the predominant type of support cell that provide structural, metabolic, visual cycle, and synaptic support (Reichenbach and Bringmann, 2013). In response to damage, certain growth factors or drug treatments can stimulate MG to become activated, de-differentiate, upregulate progenitor-related genes, re-enter the cell cycle and produce progeny that differentiate as neurons (Fischer and Bongini, 2010; Gallina et al., 2014; Wan and Goldman, 2016).

In mammalian retinas, significant stimulation such as forced expression of *Ascl1*, inhibition of histone deacetylases and neuronal damage is required to reprogram MG into progenitor-like cells that produce a few new neurons (Jorstad et al., 2017; Pollak et al., 2013; Ueki et al., 2015). Alternatively, deletion of *Nfia, Nfib* and *Nfix* in mature MG combined with retinal damage and treatment with insulin+FGF2 results in reprogramming of MG into cells that resemble inner retinal neurons (Hoang et al., 2020). In the chick retina, MG readily reprogram into progenitor-like cells that proliferate, but the progeny have a limited capacity to differentiate as neurons (Fischer and Reh, 2001a; Fischer and Reh, 2003a). Understanding the mechanisms that regulate the formation of MGPCs and neuronal differentiation of progeny is important to harnessing the regenerative potential of MG in warm-blooded vertebrates.

The responses of immune cells to damage have a profound impact upon the ability of MG to reprogram into progenitor-like in different model systems. In zebrafish, microglia influence the ability of MG to regenerate retinal neurons; the absence of reactive microglia slows the process of neuronal regeneration (Huang et al., 2012; White et al., 2017). In the mouse, the ablation of microglia enhances the generation of neuron-like cells from Ascl1-overexpressing MG in the retina (Todd et al., 2020). Reactive microglia rapidly upregulate pro-inflammatory cytokines to damaged retinas (Todd et al., 2019), and these cytokines activate NFkB-signaling in MG which, in turn, suppresses the neurogenic potential of Ascl1-overexpressing MG (Palazzo et al., 2022). In the chick, the ablation of microglia from the retina blocks the formation of proliferating MGPCs in damaged chick retinas (Fischer et al., 2014). Further, we find that the impact of NFkB-signaling on the formation of MGPCs is reversed in the absence of reactive microglia in damaged chick retinas; activation of NFkB suppresses the MGPC formation whereas in the absence of microglia activation of NFkB starts the process (Palazzo et al., 2020). Accordingly, the purpose of this study was to investigate the mechanisms underlying the failure of MGPC formation in chick retinas when the microglia/macrophage have been ablated.

## Methods and Materials

### Animals

The animals approved for use in these experiments was in accordance with the guidelines established by the National Institutes of Health and IACUC at The Ohio State University. Newly hatched P0 wildtype leghorn chicks (*Gallus gallus domesticus*) were obtained from Meyer Hatchery (Polk, Ohio). Post-hatch chicks were maintained in a regular diurnal cycle of 12 hours light, 12 hours dark (8:00 AM-8:00 PM). Chicks were housed in stainless-steel brooders at 25°C and received water and Purina^tm^ chick starter *ad libitum*.

### Intraocular injections

Chicks were anesthetized with 2.5% isoflurane mixed with oxygen from a non-rebreathing vaporizer. The technical procedures for intraocular injections were performed as previously described (Fischer et al., 1998). With all injection paradigms, both pharmacological and vehicle treatments were administered to the right and left eye respectively. Compounds were injected in 20 μl sterile saline with 0.05 mg/ml bovine serum albumin added as a carrier. Compounds included: NMDA (38.5nmol or 154 µg/dose; Millipore Sigma), FGF2 (250 ng/dose; R&D systems), BMS309403 (Millipore Sigma), C75 (Millipore Sigma), G28UCM (Millipore Sigma). 5-Ethynyl-2’-deoxyuridine (EdU, ThermoFisher) was injected into the vitreous chamber to label proliferating cells. Injection paradigms are included in each figure.

### Single Cell RNA sequencing

Retinas were obtained from embryonic and hatched chicks. Retinas were dissociated in a 0.25% papain solution in Hank’s balanced salt solution (HBSS), pH = 7.4, for 30 minutes, and suspensions were frequently triturated. The dissociated cells were passed through a sterile 70µm filter to remove large particulate debris. Dissociated cells were assessed for viability (Countess II; Invitrogen) and cell-density diluted to 700 cell/µl. Each single cell cDNA library was prepared for a target of 10,000 cells per sample. The cell suspension and Chromium Single Cell 3’ V2 or V3 reagents (10X Genomics) were loaded onto chips to capture individual cells with individual gel beads in emulsion (GEMs) using the 10X Chromium Cell Controller. cDNA and library amplification and for an optimal signal was 12 and 10 cycles respectively. Sequencing was conducted on Illumina HiSeq2500 (Genomics Resource Core Facility, John’s Hopkins University), or Novaseq6000 (Novogene) using 150 paired-end reads. Fasta sequence files were de-multiplexed, aligned, and annotated using the chick ENSMBL database (GRCg6a, Ensembl release 94) by using 10X Cell Ranger software. Gene expression was counted using unique molecular identifier bar codes and gene-cell matrices were constructed. Using Seurat toolkits Uniform Manifold Approximation and Projection for Dimension Reduction (UMAP) plots were generated from aggregates of multiple scRNA-seq libraries (Butler et al., 2018; Satija et al., 2015). Seurat was used to construct gene lists for differentially expressed genes (DEGs), violin/scatter plots and dot plots. Significance of difference in violin/scatter plots was determined using a Wilcoxon Rank Sum test with Bonferroni correction. SingleCellSignalR was used to assess potential ligand-receptor interactions between cells within scRNA-seq datasets (Cabello-Aguilar et al., 2020). Ligand-receptor interaction networks were visualized using Cytoscape (Shannon et al., 2003). Genes that were used to identify different types of retinal cells included the following: (1) Müller glia: *GLUL, VIM, SCL1A3, RLBP1*, (2) MGPCs: *PCNA, CDK1, TOP2A, ASCL1*, (3) microglia: *C1QA, C1QB, CCL4, CSF1R, TMEM22*, (4) ganglion cells: *THY1, POU4F2, RBPMS2, NEFL, NEFM*, (5) amacrine cells: *GAD67, CALB2, TFAP2A*, (6) horizontal cells: *PROX1, CALB2, NTRK1*, (7) bipolar cells: *VSX1, OTX2, GRIK1, GABRA1*, and (7) cone photoreceptors: *CALB1, GNAT2, OPN1LW*, and (8) rod photoreceptors: *RHO, NR2E3, ARR1.* Gene Ontology (GO) enrichment analysis was performed using ShinyGO V0.72 (http://bioinformatics.sdstate.edu/go/). Some of the scRNA-seq libraries can be queried at https://proteinpaint.stjude.org/F/2019.retina.scRNA.html or gene-cell matricies downloaded from GitHub at https://github.com/fischerlab3140/ and chick retina scRNA-seq Cell Ranger output files

### Fixation, sectioning and immunocytochemistry

Retinal tissue samples were formaldehyde fixed, sectioned, and labeled via immunohistochemistry as described previously (Fischer et al., 2001; Ghai et al., 2009; Ritchey et al., 2010). Antibody dilutions and commercial sources for images used in this study are described in table 1. Observed labeling was not due to off-target labeling of secondary antibodies or tissue auto-fluorescence because sections incubated exclusively with secondary antibodies were devoid of fluorescence. Secondary antibodies utilized include donkey-anti-goat-Alexa488/568, goat-anti-rabbit-Alexa488/568/647, goat-anti-mouse-Alexa488/568/647, goat-anti-rat-Alexa488 (Life Technologies) diluted to 1:1000 in PBS and 0.2% Triton X-100.

**Table 1.** List of antibodies, working dilution, clone/catalog number and source.

| Antibody | Dilution | Host | Clone/Catalog number | Source |
| --- | --- | --- | --- | --- |
| Sox2 | 1:1000 | Goat | KOY0418121 | R&D |
| Sox9 | 1:2000 | Rabbit | AB5535 | Millipore |
| Sox10 | 1:500 | Goat | PA5-47001 | Invitrogen |
| CD45 | 1:200 | Mouse | HIS-C7 | Prionics |
| Nkx2.2 | 1:100 | Mouse | 74.5A5 | DSHB |
| PMP2 | 1:100 | Rabbit | 12717.AP | Proteintech |
| AP2-alpha | 1:50 | Mouse | AP2A | DSHB |
| Brn3a | 1:50 | Mouse | MAB1585 | Chemicon |
| Islet1 | 1:50 | Mouse | 40.2D6 | DSHB |
| Tyrosine hydroxylase | 1:50 | Mouse | aTH | DSHB |
| Olig2 | 1:50 | Mouse | PCRP-Olig2-1E9 | DSHB |
| Vimentin | 1:50 | Mouse | H5 | DSHB |
| Glutamine synthetase | 1:1000 | Mouse | 610517 | BD Biosciences |
| pS6 | 1:750 | Rabbit | #2215; Ser240/244 | Cell Signaling |
| pStat3 | 1:100 | Rabbit | 9131 | Cell Signaling |
| pSmad1/5/8 | 1:200 | Rabbit | D5B10 | Cell Signaling |

### Labeling for EdU

For the detection of nuclei that incorporated EdU, immunolabeled sections were fixed in 4% formaldehyde in 0.1M PBS pH 7.4 for 5 minutes at room temperature. Samples were washed for 5 minutes with PBS, permeabilized with 0.5% Triton X-100 in PBS for 1 minute at room temperature and washed twice for 5 minutes in PBS. Sections were incubated for 30 minutes at room temperature in a buffer consisting of 100 mM Tris, 8 mM CuSO_4_, and 100 mM ascorbic acid in dH_2_O. The Alexa Fluor 568 Azide (Thermo Fisher Scientific) was added to the buffer at a 1:100 dilution.

### Photography, measurements, cell counts and statistics

Microscopy images of retinal sections were captured with the Leica DM5000B microscope with epifluorescence and the Leica DC500 digital camera. High resolution confocal images were obtained with a Leica SP8 available in The Department of Neuroscience Imaging Facility at The Ohio State University. Representative images are modified to have enhanced color, brightness, and contrast for improved clarity using Adobe Photoshop. In EdU proliferation assays, a fixed region of retina was counted and average numbers of Sox2 and EdU co-labeled cells. The retinal region selected for investigation was standardized between treatment and control groups to reduce variability and improve reproducibility.

MS Excel and GraphPad Prism 9 were used for statistical analyses. For statistical evaluation of differences in treatments, a two-tailed paired *t*-test was applied for intra-individual variability where each biological sample also served as its own control. For two treatment groups comparing inter-individual variability, a two-tailed unpaired *t*-test was applied. For multivariate analysis, an ANOVA with the associated Tukey Test was used to evaluate any significant differences between multiple groups.

## Results

### scRNA-seq analyses of MG from retinas with and without microglia

For simplicity, hereafter, we refer to retinal microglia and infiltrating macrophage as microglia since the CD45 antibodies used to identify these cells does not distinguish between resident microglia and recruited macrophage. We generated 4 scRNA-seq libraries; (i) control undamaged retina, (ii) undamaged retinas with microglia depleted, (iii) NMDA damaged retinas at 24 hrs after treatment, and (iv) NMDA damaged retinas at 24 hrs after treatment with microglia depleted. Retinal microglia were depleted by a single intravitreal injection of clodronate-liposomes, which effectively destroys >95% of microglia/macrophage within 3 days of treatment (Fischer et al., 2014; Fischer et al., 2015; Palazzo et al., 2020; Zelinka et al., 2012). UMAP plots of aggregated libraries (43,566 total cells) revealed distinct clusters of cell types, with neuronal types clustered together regardless of treatment and MG forming distinct clusters that correlated to treatments (Fig. 1a,b). Clusters of retinal cells were identified based on cell-distinguishing markers, as described in the methods. Resting MG were identified based on patterns of expression for *GLUL, RLBP1* and *GPR37L1*, and activated MG were identified based on expression of *PMP2, TGFB2* and *MDK* (Fig. 1b-d). We bioinformatically isolated MG and re-ordered these cells in a UMAP plot. Approximately 1000 MG from each treatment group formed distinct clusters of cells that were segregated into resting and activated MG (Fig. 1c,d). We generated lists of differentially expressed genes (DEGs) that were up-or downregulated in MG in undamaged and damaged retinas, with and without microglia (Fig. 1e,g; supplemental tables 1,2,3). The depletion of microglia in undamaged retinas caused MG to significantly downregulate many (121) genes associated with resting MG, including different receptors, transcription factors, and components of different cell signaling pathways (Fig. 1g,f). Many (82) of these genes were also downregulated by MG in NMDA-damaged retinas (Fig. 1g). By comparison, the depletion of microglia in undamaged retinas caused MG to significantly upregulate many (265) genes associated with activated MG, including secreted factors, pro-inflammatory cell signaling, and transcription factors (Fig. 1e,f). We identified many DEGs in MG in undamaged retinas that resulted from the absence of microglia; downregulated DEGs included genes associated with resting glia including different receptors, transcription factors and genes associated with cell signaling pathways (Fig. 1f). Upregulated DEGs included genes involved in fatty acid metabolism and transcription factors known to promote glial phenotype (Fig. 1f). We also identified many DEGs in MG in damaged retinas that were significantly affected by the absence of microglia (Fig. 1e,g,h). Relatively few (62) upregulated DEGs were identified, whereas numerous (350) downregulated DEGs were identified in MG in damaged retinas missing microglia (Fig. 1g). Downregulated DEGs included many secreted factors known to stimulate the proliferation of MGPCs, markers of reactive glia, and genes associated with NFkB-(*TNFRSF11B, TNFRSF21, NFKBIB*), MAPK-(*FGFR1, MAPK6, DUSP1*) and Notch-signaling (*MAML2*) (Fig. 1g,h). Among the most significantly downregulated genes in MG in damaged retinas with microglia depleted were secreted factors known to stimulate the formation of proliferating MGPCs, including Wnt ligands (Gallina et al., 2015), HBEGF (Todd et al., 2015) and midkine (Campbell et al., 2021a).

**Figure 1.**
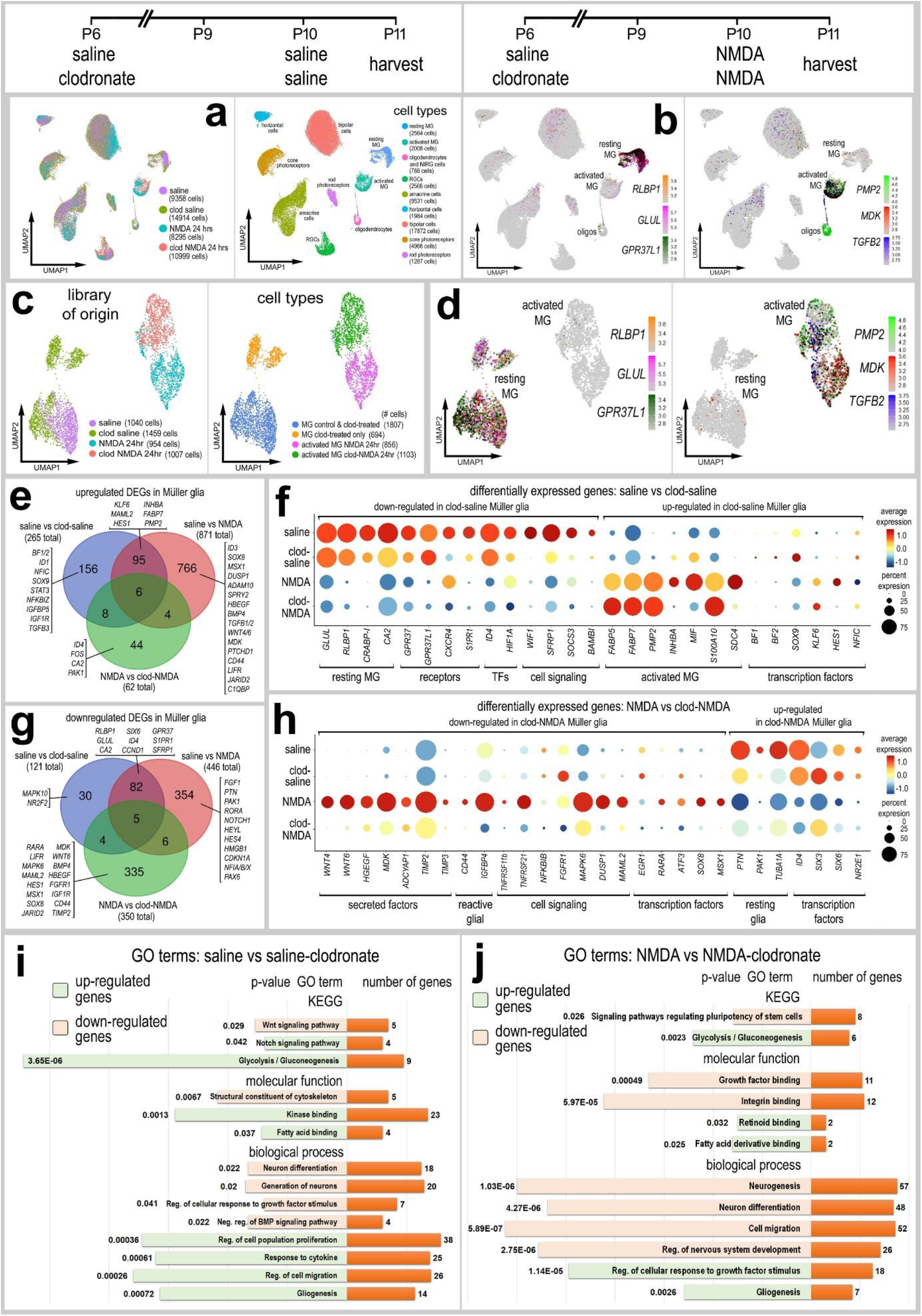
scRNA-seq of normal and damaged retinas with and without microglia. Retinas were obtained from eyes injected with saline or clodronate liposomes at P6, saline or NMDA at P10, and tissues harvested at 24hrs after the last injection. UMAP ordering of cells is displayed for libraries of origin and distinct clusters of cells (**a**). In UMAP heatmap plots, resting MG were identified based on elevated expression of *RLBP1, GLUL* and *GPR37L1* (**b,d**), and activated MG were identified based on elevated expression of *PMP2, MDK* and *TGFB2* (**b,d**). MG were bioinformatically isolated and re-embedded in UMAP plots (**c,d**). Lists of DEGs were generated (supplemental tables 1,2,3) for MG from retinas treated with saline vs clodronate-saline, saline vs NMDA, and NMDA vs clodronate-NMDA. Numbers of up- and down-regulated DEGs are plotted in a Venn diagrams (**e,g**). Genes of interest unique to each treatment are listed. Dot plots illustrate the percentage of expressing MG (size) and significant (p<0.01) changes in expression levels (heatmap) for genes in MG from retinas treated with saline vs saline+ clodronate (**f**) and NMDA vs NMDA+clodronate (**h**). Significance of difference was determined by using a Wilcox rank sum with Bonferoni correction. GO enrichment analysis was performed for lists of DEGs in MG for retinas treated with saline ± clodronate and NMDA ± clodronate (**i,j**). Gene modules for upregulated (green) and downregulated (peach) genes were grouped by GO category with P values and numbers of genes for each category (**i,j**).

We performed Gene Ontology (GO) enrichment analyses of lists of up- and downregulated genes in MG in undamaged retinas without microglia. Depletion of microglia from undamaged retinas stimulated MG to downregulate gene modules associated with Wnt-signaling, BMP-signaling, and neurogenesis (Fig.1i). By comparison, depletion of microglia from undamaged retinas stimulated MG to upregulate gene modules associated with Notch-signaling, fatty acid binding, glycolysis, cellular proliferation/migration and gliogenesis (Fig. 1i). GO enrichment analyses of lists of up- and downregulated genes in MG in damaged retinas without microglia revealed upregulated gene modules associated with glycolysis, fatty acid binding, and gliogenesis, similar to gene modules identified in undamaged retinas missing microglia (Fig. 1j). By comparison, depletion of microglia from damaged retinas stimulated MG to downregulate gene modules associated with regulation of stem cell pluripotency, growth factor/integrin binding and neuronal development/differentiation (Fig. 1j).

### Implied Ligand-Receptor interactions that change when microglia are ablated

We next bioinformatically isolated the MG and re-embedded these cells to probe for cell signaling networks and putative ligand-receptor (LR) interactions using SingleCellSignalR (Cabello-Aguilar et al., 2020). We started by analyzing autocrine signaling among MG that might be affect by the ablation of microglia/macrophage. Resting MG included cells for saline- and clodronate-treatment groups, activated MG included cells from NMDA and clodronate-NMDA treatment groups. Numbers of LR-interactions (significant upregulation of putative ligand and receptor) between cell types in the different treatment groups varied between 70 and 315. We performed analyses on MG from each treatment group and compared changes across the most significant LR-interactions (Fig. 2a-d). Among resting MG LR-interactions included signaling through different intergrins, FGFR4, and pleotrophin (PTN)-PTPRZ1 (Fig. 2a,b,e). By comparison, depletion of microglia included signaling through different ligand-integrin pairs, midkine (MDK)/SDC1/4, EGF/EGFR and FGF18/FGFR1 (Fig. 2a,b,f). Among activated MG in damaged retinas, we observed significant LR-interactions involving HBEGF/EGFR, FGF18/FGFR1, and SMAD3 (Fig. 2c,d,f). When the microglia were ablated (Fig. 2a,e). By comparison, MG with microglia ablated included LR-interactions with EGFR and FGFR and interactions involving integrins ITGB1 and ITGAV (Fig. 2b,e).

**Figure 2.**
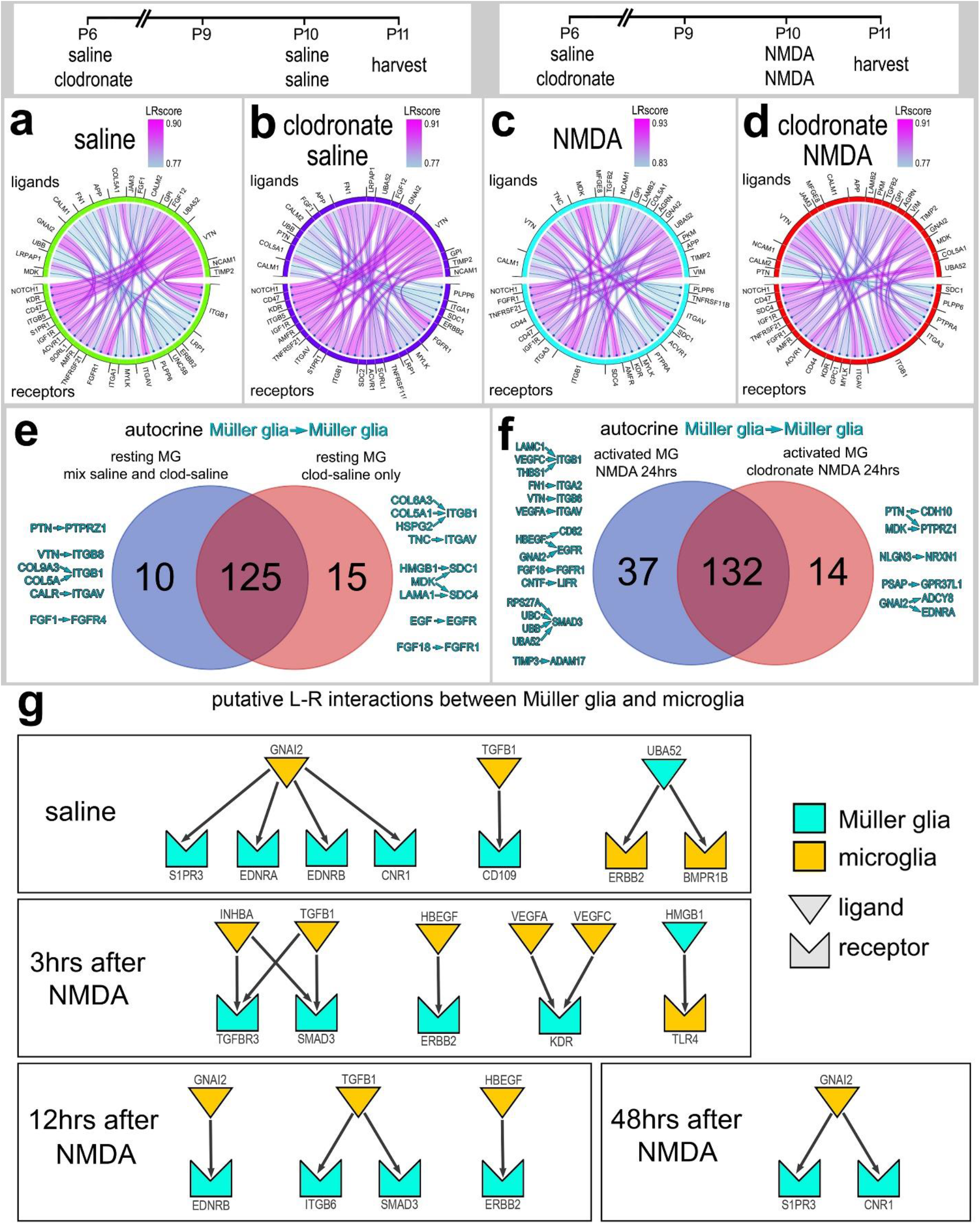
Inferred autocrine ligand-receptor (LR) interactions between MG. SingleCellSignalR was used to identify putative LR interactions. Chord diagrams illustrate the top 30 most significant autocrine LR interactions between MG in retinas treated with saline (**a**), clodronate-saline (**b**), NMDA (**c**) and clodronate-NMDA (**d**). Significant autocrine interactions were compared for saline vs clodronate-saline and NMDA vs clodronate-NMDA. Venn diagrams illustrate numbers of unique and common LR interactions between treatment groups (**e,f**). Representative unique LR-interactions for undamaged retinas with and without microglia (**e**) and NMDA damaged retinas with and without microglia (**f**). Representative significant LR-interactions between microglia and MG are illustrated for undamaged and damaged retinas with or without microglia (**g**).

We bioinformatically isolated MG and microglia from normal and NMDA-damaged retinas at 3, 12 and 48hrs after treatment to identify putative LR-interactions. (Fig. 2g). We identified LR-interactions between microglia and MG in undamaged retinas involving ligands (CNR1, TGFB1) and receptors (ERBB2, BMPR1B) (Fig. 2g, supplemental Fig. 1) known to influence the reprogramming of MG (Campbell et al., 2021b; Todd et al., 2015; Todd et al., 2017). In NMDA-damaged retinas, we identified LR-interactions between microglia and MG involving TGFB1/TGFBR3/SMAD3 and HBEGF/ERBB2 (Fig. 2g, supplemental Fig. 1), pathways known to inhibit or activate, respectively, the formation of MGPCs (Todd et al., 2015; Todd et al., 2017). A recent study in zebrafish retinas provides evidence that expression of Vegfa is induced in MG by reactive microglia/macrophages to promote the reprogramming of MG into proliferating MGPCs (Mitra et al., 2022).

### Wnt signaling and the formation of MGPCs in damaged retinas missing microglia

Wnt-ligands, *WNT4* and *WNT6*, are among the most significantly downregulated genes in MG in damaged retinas where microglia were ablated. Activation of Wnt-signaling is known to stimulate the formation of proliferating MGPCs in the retinas of fish (Ramachandran et al., 2011), chicks (Gallina et al., 2015), and rodents (Osakada et al., 2007; Yao et al., 2016). We have previously reported that intraocular injections of Wnt agonists or a cocktail of GSK3β-inhibitors stimulated the proliferation of MGPCs in damaged chick retinas (Gallina et al., 2016). Accordingly, we probed for expression levels of Wnt-related genes in scRNA-seq libraries of retinas obtained at 3, 12 and 48 hrs after NMDA (Fig. 3a).

**Figure 3.**
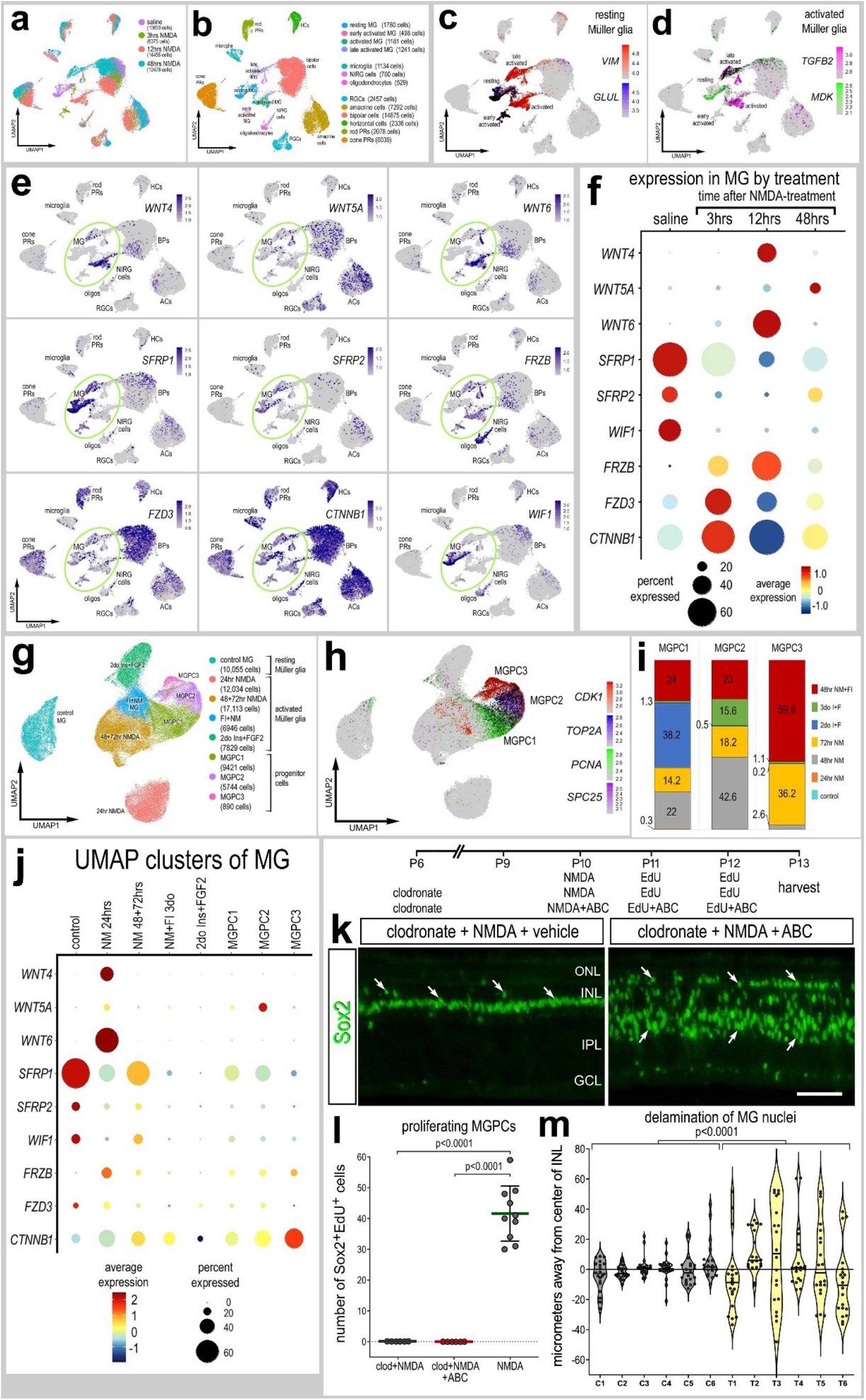
Patterns of expression of Wnt-related genes in normal and damaged retinas. scRNA-seq was used to identify patterns and levels of expression of Wnt-related genes in control and NMDA-damaged retinas at 3, 12 and 48 hours after treatment (**a**). UMAP clusters of cells were identified based on well-established patterns of gene expression (see Methods; **b**). MG were identified by expression of *VIM* and *GLUL* in resting MG (**c**), and *TGFB2* and *MDK* activated MG (**d**). Each dot represents one cell and black dots indicate cells that express 2 or more genes (**c,d**). UMAP heatmap plots (feature plots) (**e**) illustrate patterns and levels of expression for Wnt-related genes in retinal neurons and glia. The dot plots in **f** and **j** illustrate average expression (heatmap) and percent expressed (dot size) for different Wnt-related genes in MG that are significantly (p<0.01) up- or downregulated at different times following NMDA-treatment. Significance of difference was determined by using a Wilcox rank sum test with Bonferroni correction. MG were bioinformatically isolated from scRNA-seq libraries of control retinas, retinas at 24, 48 and 72hrs times after NMDA treatment, retinas treated with 2 or 3 doses of insulin and FGF2, and retinas treated with insulin, FGF2 and NMDA at 48hrs after NMDA for a total of 70,032 MG (**g-j**). MG formed distinct UMAP clusters that correlated to different treatment and MGPCs for clusters based on progression through the cell cycle (**h**). Clusters of MGPCs were occupied by cells from different treatment groups (**i**). **Inhibition of GSK3β to stimulate Wnt-signaling fails to stimulate MGPCs proliferation, but induces delamination of MG nuclei**. We applied clodronate liposomes at P6, NMDA a cocktail of GSK3β inhibitors at P10, EdU GSK3β inhibitors at P11 and P12, and harvested retinas at P13. Retinal sections were labeled for EdU accumulation and antibodies to Sox2 (green; **k**). **(l)** Histogram representing mean MGPC proliferation (bar ± SD); significance of difference (p-values) was determined using an unpaired t-test. Each dot represents one biological replicate. (**m**) Violin plot representing MG nuclear delamination; significance of difference (p-values) was determined using one-way ANOVA. Each dot represents one biological replicate. Calibration bar in **k** represent 50 µm. Arrows indicate the nuclei of MG. Abbreviations: ONL – outer nuclear layer, INL – inner nuclear layer, IPL – inner plexiform layer, GCL – ganglion cell layer.

After filtering of cells with low numbers of genes/cells, putative doublets (high numbers of genes/cells) and dying cells (high percentage of mitochondrial genes), we analyzed a total of 42,202 cells including a total of 4,700 MG (Fig. 3a,b). UMAP ordering of cells formed discrete clusters of neurons according to cell types and MG according to treatment (Fig.3a,b). MG were identified based on expression of *VIM*, resting MG (*GLUL*) and activated MG were identified based on expression of *MDK* and *TGFB2* (Fig. 3c,d). *WNT4* and *WNT6* were expressed at low levels by resting MG and activated MG at 3 and 48hrs after NMDA, but were highly expressed by activated MG at 12hrs after NMDA (Fig. 3e,f). By comparison, *WNT5A* was upregulated by MG at 48hrs, but was also expressed by some retinal ganglion, amacrine and bipolar cells (Fig. 3e,f). Wnt inhibitors *SFRP1, SFRP2* and *WIF1* were highly expressed by resting MG and rapidly downregulated by 3hrs after NMDA and remained low through 48hrs after treatment (Fig. 3e,f). By contrast, the Wnt inhibitor *FRZB* was low in resting MG and elevated at 3 and 12hrs after NMDA, and was highly expressed by oligodendrocytes and NIRG cells (Fig. 3e,f). *Wnt* receptor FZD3 and transcriptional effector *CTNNB1* (β-catenin) were rapidly upregulated at 3 hours, downregulated at 12hrs, and back up at 48hrs after NMDA-treatment (Fig. 3e,f). Collectively, these findings suggest Wnt-related genes are rapidly (3hrs or less) and transiently up- or down-regulated in MG following damage to retinal neurons.

To gain a more comprehensive understanding of patterns of expression of Wnt-related genes, we probed a large aggregate scRNA-seq library of MG (>70,000 cells) with glia bioinformatically isolated from retinas treated with saline, 24/48/72hrs NMDA, 2 or 3 doses of insulin+FGF2 and 48hrs NMDA+insulin+FGF2 (Fig. 3g,h), as described previously (Campbell et al., 2021a; Campbell et al., 2021b; El-Hodiri et al., 2022). Resting MG and MG at 24hrs after NMDA formed 2 distinct clusters of cells, whereas activated MG from different treatment groups and MGPCs formed a continuum of cells based, in part, on expression of cell cycle regulators (Fig. 3g-j). *WNT4* and *WNT6* were exclusively upregulated by the MG from 24hrs after NMDA-treatment (Fig. 3j). *WNT5A* was most highly expressed in the MGPC1 cluster which was predominantly occupied by MG from retinal libraries from 48 and 72hrs after NMDA, 2 doses of insulin+FGF2, and 48hrs after NMDA+insulin+FGF2 (Fig. 3i,j). Wnt inhibitors *SRFP1, SFRP2* and *WIF1* were highest in resting glia and reduced in activated MG and MGPCs from all treatments (Fig. 3j). *FRZB* was highest in MG at 24hrs after NMDA and was relatively decreased in all other MG (Fig. 3j). Wnt-receptor *FZD3* was relatively high in resting MG and decreased in all other treatment groups, except for the MG at 3hrs after NMDA represented in the other aggregate library (Fig. 3f,j). *CTNNB1* (β-catenin) was most highly expressed in the MGPC3 cluster which was predominantly occupied by MG from retinal libraries from 72 hrs after NMDA and 48hrs after NMDA+insulin+FGF2 (Fig. 3i,j). Collectively, these findings suggest Wnt-related genes are transiently up- or downregulated in MG following treatments that stimulate the formation of MGPCs, regardless of retinal damage.

We next tested whether intraocular injections of GSK3β-inhibitors rescued the deficit of MGPC proliferation in damaged retinas missing reactive microglia. The biological activity of recombinant Wnt-ligands remains uncertain because these proteins do not have correct conformational folding that is necessary for biological function. Indeed, we found that injections of recombinant Wnt4 failed to stimulate the nuclear localization of β-catenin and formation of proliferating MGPCs in damaged retinas missing microglia (not shown). We have previously shown that nuclear localization of β-catenin is maximal in proliferating MGPCs at 2-3 days after NMDA-treatment (REF Gallina et al., 2016). Accordingly, we injected a cocktail of GSK3β inhibitors which have been shown to stimulate the accumulation of nuclear β-catenin and the proliferation of MGPCs in damaged retinas (Gallina et al., 2015). We found GSK3β inhibitors failed to stimulate the proliferation of MGPCs, but did cause widespread delamination of MG nuclei (Fig. 3k-m). In all treated individuals, we observed a significant migration of Sox2-positive MG nuclei away from the middle of the INL (Fig. 3k,m).

### Exogenous FGF2 rescues the failure of MGPC proliferation in damaged retinas missing microglia

Analyses of scRNA-seq libraries of MG from undamaged and NMDA-damaged retinas with and without microglia indicated that components of FGF/MAPK-signaling were upregulated by MG in damaged retinas and this upregulation is diminished when microglia are absent (Fig. 1a). These components included *FGFR1, MAPK6* and *SPRY2* (Fig. 1a). Interestingly, *FGFR1* was significantly upregulated in MG in undamaged retinas when microglia were ablated (Fig.1a). Other MAPK-related factors that were up- or downregulated with NMDA-treatment were not significantly affected by the ablation of microglia.

Consistent with previous reports, a single intravitreal injection of clodronate-liposomes effectively eliminated microglia from the retina (Fig.4b). We tested whether addition of exogenous FGF2 rescued the deficit in MGPC proliferation in damaged retinas missing microglia. Injections of FGF2 with and following NMDA significantly increased numbers of proliferating MGPCs in retinas missing microglia (Fig. 4c,d). The number of proliferating MGPCs in clodronate/NMDA/FGF2-treated retinas was not significantly different from numbers seen in retinas treated with NMDA alone (Fig. 4d), suggesting a complete rescue.

**Figure 4.**
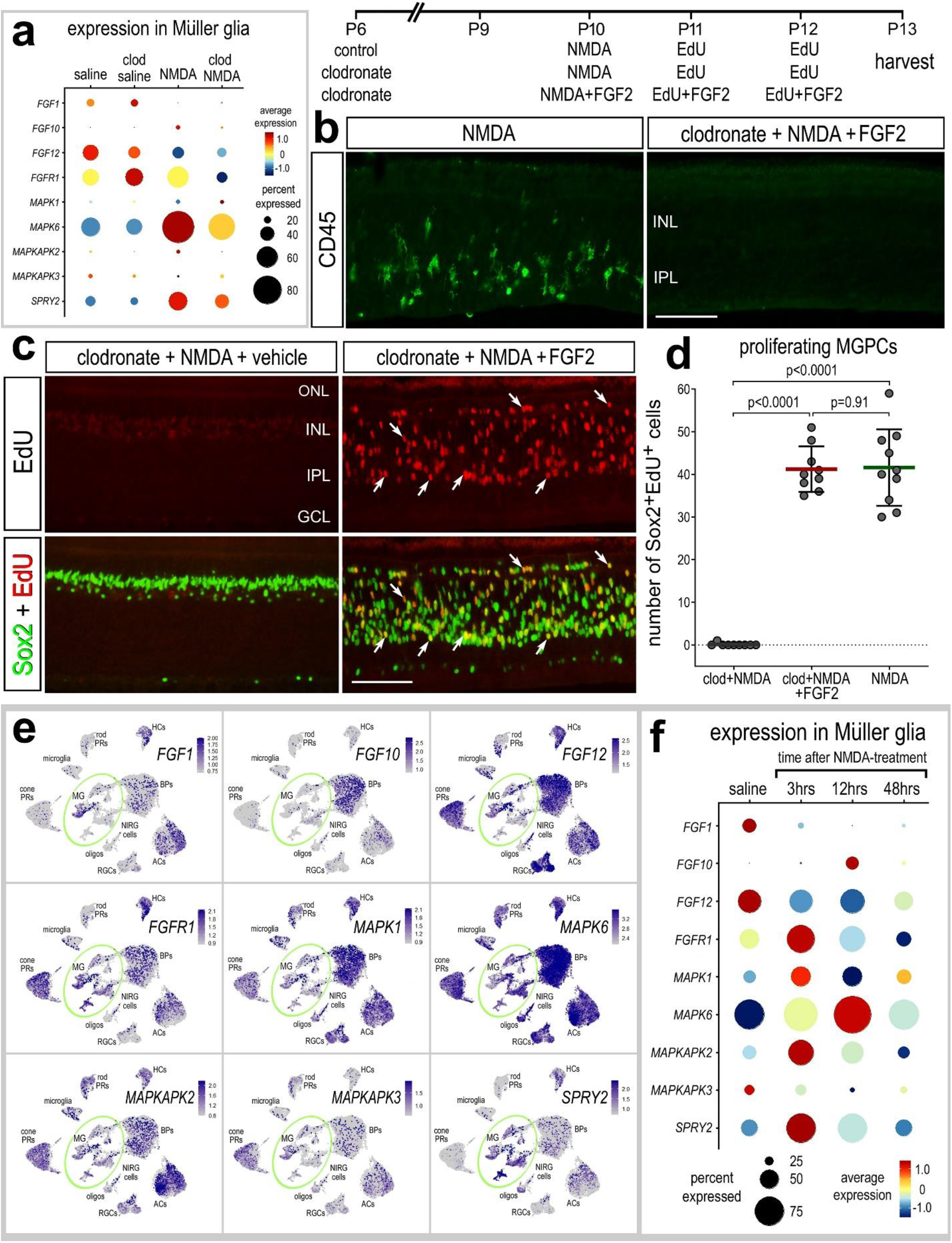
Activation of FGF2 rescues the deficit in MGPC proliferation in damaged retinas where microglia have been ablated. scRNA-seq was used to analyze patterns and levels of expression of FGF/MAPK-related genes in retinas treated with saline ± clodronate and NMDA ± clodronate (**a**), or different times after NMDA treatment (**e,f**). The dot plots in **a** and **f** illustrate expression levels (heatmap) and percent expressed (dot size) for different genes in MG that are significantly up- or downregulated. Significance of difference (p<0.01) was determined by using a Wilcox rank sum test with Bonferroni correction. Retinal sections were labeled for EdU accumulation (red; **c**) and/or antibodies to Sox2 (green; **c**) and CD45 (green; **b**). The histogram in **d** illustrates the mean (bar ± SD) and each dot represents one biological replicate. Significance of difference (p-values) was determined by using an unpaired t-test with bonferroni correction (**d**). Calibration bars in **b** and **c** represent 50 µm. Arrows indicate the nuclei of MG. UMAP heatmap plots (**e**) illustrate patterns and levels of expression for FGF/MAPK-related genes in retinal neurons and glia.

To better understand the time-course of changes in expression, we probed for expression levels of FGF/MAPK-related genes in the retina and MG in scRNA-seq libraries generated shortly (3 or 12 hrs) after NMDA-treatment. *FGF1, FGF12* and *MAPKAPK3* were rapidly (<3hrs) downregulated by MG following NMDA-treatment (Fig. 4e,f). By contrast, *FGF10, FGFR1, MAPK1, MAPK6, MAPKAPK2* and *SPRY2* were rapidly (3-12 hrs) upregulated by MG following NMDA-treatment (Fig. 4e-f). These genes were downregulated by 48hrs after NMDA-treatment (Fig. 4f). Collectively, these findings suggest that components of MAPK-signaling are very quickly modulated in MG following neuronal damage.

### Exogenous HBEGF rescues the failure of MGPC proliferation in damaged retinas missing microglia

MG significantly upregulate HBEGF after NMDA, and injections of HBEGF stimulate the proliferation of MGPCs (Todd et al., 2015) and Fig. 1h). We found that the upregulation of *HBEGF* by MG was greatly diminished when microglia were absent (Fig. 1h, 5a,b). *EGFR* was not expressed by MG, but there was a small, but significant increase in *ERBB2* in MG in damaged retinas, and this increase was absent when microglia where ablated (Fig. 5a,b). By comparison, *GRB2* was expressed at relatively high levels in resting MG, decreased in activated MG, and levels were decreased under both conditions when microglia were ablated (Fig. 5a,b). We next probed for patterns of expression of *ADAM9* and *ADAM10*, enzymes known to process HBEGF in the extracellular space. ADAM9 is involved in Tissue Plasminogen Activator (TPA) -induced ectodomain shedding of membrane tethered HBEGF. ADAM10 proteolytically releases cell-surface proteins, including HBEGF, ephrin-A2, CD44 and CDH2 (Jouannet et al., 2016; Lemjabbar and Basbaum, 2002; Seegar et al., 2017). *ADAM9* was expressed by resting MG and was downregulated at 24hrs after NMDA-treatment and when the microglia were ablated in undamaged retinas, but was upregulated by MG at 24hrs after NMDA when microglia were ablated (Fig. 5a,b). Similar to patterns of expression of *HBEGF*, *ADAM10* was upregulated by MG at 24hrs after NMDA, and this upregulation was diminished when microglia were ablated (Fig. 5a,b).

**Figure 5.**
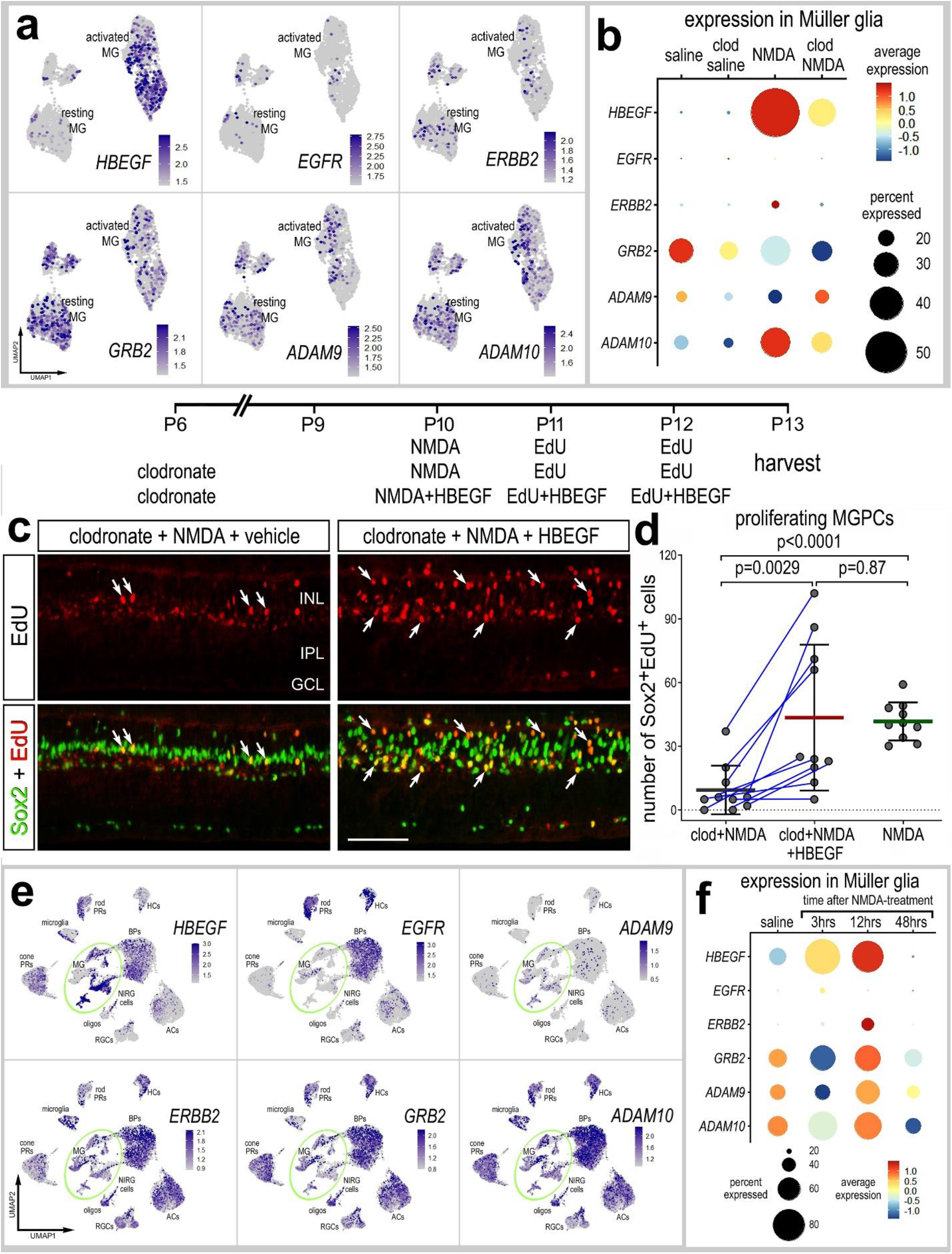
Activation of HBEGF rescues the deficit in MGPC formation in damaged retinas where microglia have been ablated. scRNA-seq was used to analyze patterns and levels of expression of EGF-related genes in retinas treated with saline ± clodronate and NMDA ± clodronate (**a,b**), or different times after NMDA treatment (**e,f**). UMAP heatmap plots (feature plots) illustrate patterns and levels of expression for EGF-related genes in MG only (**a**) or across all types of retinal neurons and glia (**e**). The dot plots in **b** and **f** illustrate expression levels (heatmap) and percent expressed (dot size) for different genes in MG that are significantly up- or downregulated. Significance of difference (p<0.01) was determined by using a Wilcox rank sum test with Bonferroni correction. Retinal sections were labeled for EdU accumulation (red; **c**) and antibodies to Sox2 (green; **c**). The histogram in **d** illustrates the mean (bar ± SD) and each dot represents one biological replicate. Significance of difference (p-values) was determined by using an unpaired t-test with Bonferroni correction (**d**). Calibration bars in **c** represent 50 µm. Arrows indicate the nuclei of MG.

We tested whether addition of exogenous HBEGF rescued the deficit of MGPC proliferation in damaged retinas missing reactive microglia. HBEGF is known to stimulate the formation of proliferating MGPCs in the retinas of fish (Wan et al., 2012), chicks and mice (Todd et al., 2015). We found that injections of HBEGF with and following NMDA significantly increased numbers of proliferating MGPCs in retinas missing microglia (Fig. 5c,d). Consistent with this observation, we found that the Sox2+ nuclei of MG migrated away from the center of the INL (Fig. 5c,d), consistent with previous reports wherein delamination of MG nuclei is part of the process of reprogramming into MGPCs.

To better understand the time-course of changes in expression, we probed for expression levels of EGF-related genes in the retina and MG in scRNA-seq libraries generated shortly (3 or 12 hrs) after NMDA-treatment. We found that *HBEGF* was rapidly upregulated within 3hrs of NMDA-treatment (Fig. 5e,f). Levels of *HBEGF* were further increased at 12hrs and were absent by 48hrs after NMDA-treatment (Fig. 5e,f). We next probed for expression of EGF receptors which include *ERBB2, EGFR* and *GRB2*. *ERBB2* was not widely expressed by different types of retinal cells (Fig. 5e), whereas *EGFR* was highly expressed by bipolar cells, rod photoreceptors and horizontal cells (Fig. 5e). *ADAM9* was not widely expressed by different types of retinal neurons, but was expressed by resting MG and was downregulated at 3hrs, up at 12hrs and back down by 48hrs after NMDA treatment (Fig. 5e,f). By comparison, *ADAM10* was widely expressed by different types of retinal neurons and glia (Fig. 5e). Similar to patterns of expression of *ADAM9* in MG, *ADAM10* was downregulated at 3hrs, up at 12hrs and back down by 48hrs after NMDA treatment (Fig. 5f).

### Inhibition of Smad3 and the formation of MGPCs in in the absence of microglia

Signaling through TGFβ2 and Smad2/3 has been shown to act, in opposition to BMP and Smad1/5/8, to suppress the formation of proliferating MGPCs (Todd et al., 2017). Accordingly, we probed for changes in expression levels of TGFβ-related genes in MG in normal and damaged retinas, with and without retinas. We found the absence of microglia from damaged retinas, results in elevated levels of *TGFB1, TGFB2* and *TGFBR1* and decreased levels of inhibitors *TGIF1* and *INHBA* (Fig. 6a,b). We tested whether inhibition of TGFβ/Smad-signaling rescued the deficit in MGPC proliferation in damaged retinas missing microglia. Injections of Smad3-antagonist (SIS3) to NMDA-damaged retinas missing microglia resulted in a small, but significant increase in numbers of proliferating MGPCs (Fig. 6c,d). This increase in MGPC proliferation was significantly less that that seen in retinas treated with NMDA alone (Fig. 6d).

**Figure 6.**
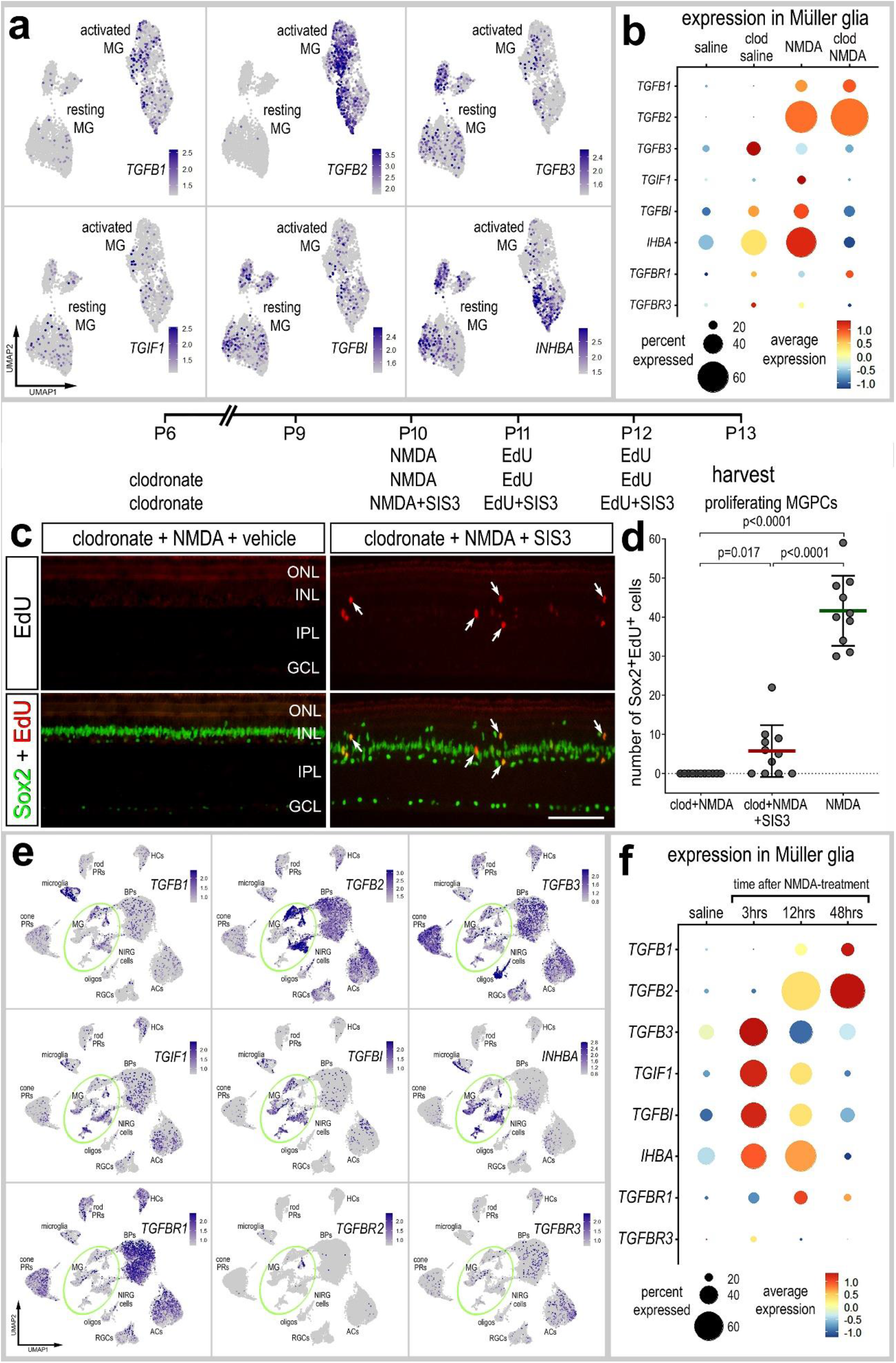
Activation of TGFβ/Smad3-signaling partially rescues the deficit in MGPC formation in damaged retinas where microglia have been ablated. scRNA-seq was used to analyze patterns and levels of expression of TGFβ-related genes in retinas treated with saline ± clodronate and NMDA ± clodronate (**a,b**), or different times after NMDA treatment (**e,f**). UMAP heatmap plots illustrate patterns and levels of expression for TGFβ-related genes in MG only (**a**) or across all types of retinal neurons and glia (**e**). The dot plots in **b** and **f** illustrate expression levels (heatmap) and percent expressed (dot size) for different genes in MG that are significantly up- or downregulated. Significance of difference (p<0.01) was determined by using a Wilcox rank sum test with Bonferroni correction. Retinal sections were labeled for EdU accumulation (red; **c**) and antibodies to Sox2 (green; **c**). The histogram in **d** illustrates the mean (bar ± SD) and each dot represents one biological replicate. Significance of difference (p-values) was determined by using an unpaired t-test with Bonferroni correction (**d**). Calibration bars in **c** represent 50 µm. Arrows indicate the nuclei of MG.

To better understand the time-course of changes in expression, we probed for expression levels of TGFβ-related genes in the retina and MG in scRNA-seq libraries generated shortly (3 or 12 hrs) after NMDA-treatment. *TGFB1* was predominantly expressed by microglia (Fig. 6e). By comparison, *TGFB2* was predominantly expressed by activated MG, bipolar cells and amacrine cells (Fig. 6e,f). *TGFB3* was widely expressed by cone photoreceptors and oligodendrocytes, and showed scattered expression across all othe types of retina neurons and glia (Fig. 6e.). In MG, specifically, *TGFB1* and *TGFB2* were significantly upregulated at 12 and 48hrs after NMDA, with *TGFB2* being expressed by most MG (Fig. 6f), whereas *TGFB3* was significantly upregulated in MG at 3hrs after NMDA and downregulated thereafter (Fig. 6f). We next probed for genes related to TGFβ-signaling, including *TGIF1, TGFBI* and I*NHBA*. *TGIF1* (TGFβ Induced Factor Homeobox 1) is a transcriptional co-repressor of Smad2 to regulate TGFβ-signaling (Guca et al., 2018). TGFBI (Transforming Growth Factor Beta Induced) is induced by TGFβ and acts to inhibit cell adhesion (LeBaron et al., 1995; Skonier et al., 1994). *INHBA* encodes a member of the TGFβ superfamily of proteins; the preproprotein is proteolytically processed to generate a subunit of the dimeric activin and inhibin protein complexes (Antenos et al., 2008; Ling et al., 1986). *TGIF1, TGFBI* and *INHBA* were predominantly expressed in MG (Fig. 6e,f). Similar to patterns of expression of *TGFB3*, levels of these factors were low in resting MG, upregulated within 3hrs, remained elevated at 12hrs, and downregulated by 48hrs after NMDA-treatment (Fig. 6f). TGFβ-receptors, *TGFBR1* and *TGFBR3,* were expressed by different types of retinal neurons and glia, whereas *TGFBR2* was not widely expressed in the retina (Fig. 6e). *TGFBR1* was upregulated by a relatively small percentage of MG at 12hrs after treatment, whereas *TGFBR3* was not expressed by MG (Fig. 6f).

### Activation of RARα and the formation of MGPCs in retinas missing microglia

Signaling through retinoic acid receptors is known to stimulate the proliferation of MGPCs and increase numbers of progeny that differentiate as neurons (Todd et al., 2018). One of the key receptors of retinoic acid (RA) is *RARA*, which is highly upregulated by MG in damaged retinas, but fails to become upregulated when microglia are absent (Fig. 1g,h, Fig. 7a,b). By comparison, *RARB*, *RXRA* and *RXRG* are downregulated by MG in damaged retinas, but are upregulated in resting MG when microglia are ablated (Fig. 7a,b). Similarly, *CYP26A1*, an enzyme that inactivates RA via oxidation, and *ALDH1A1*, an enzyme involved in the synthesis of RA, are down regulated in activated MG in damaged retinas (Fig. 7a,b). *ALDH1A2* was not detected at significant levels in the chick retina (not shown). Injections of RAR-agonist, TTNBP, to NMDA-damaged retinas missing microglia resulted in a small, but significant increase in numbers of proliferating MGPCs labeled for EdU and Sox2 (Fig. 7c,d). Worth noting, intraocular injections of TTNBP compromised the survival of about 50% the chicks; we have not observed this effect for dozens of other compounds that we have applied to the eye of chicks over the past 25 years.

**Figure 7.**
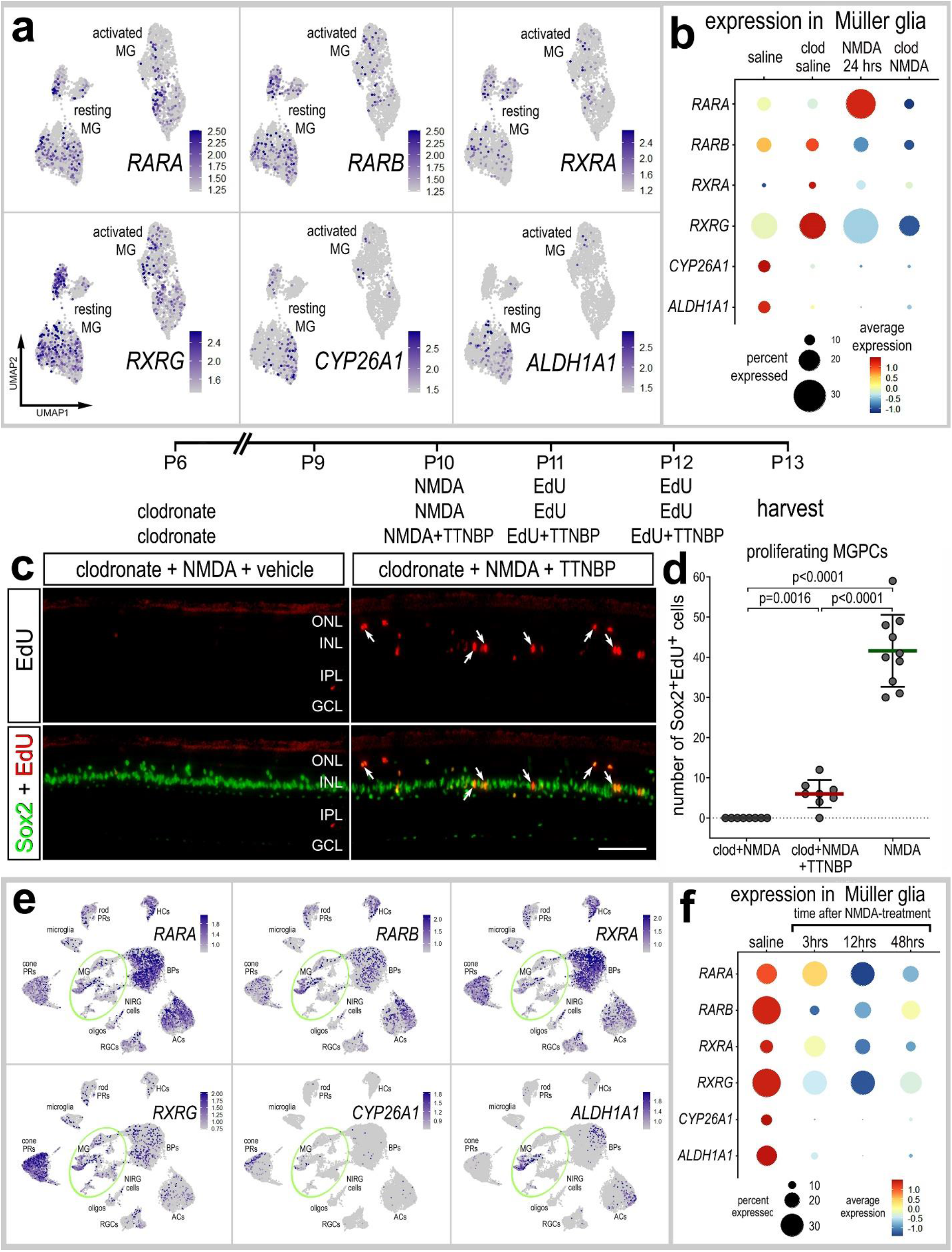
Inhibition of retinoic acid-signaling partially rescues the deficit in MGPC formation in damaged retinas where microglia have been ablated. scRNA-seq was used to analyze patterns and levels of expression of retinoic acid-related genes in retinas treated with saline ± clodronate and NMDA ± clodronate (**a,b**), or different times after NMDA (**e,f**). UMAP heatmap plots (feature plots) illustrate patterns and levels of expression for retinoic acid-related genes in MG only (**a**) or across all types of retinal neurons and glia (**e**). The dot plots in **b** and **f** illustrate expression levels (heatmap) and percent expressed (dot size) for different genes in MG that are significantly up- or downregulated. Significant of difference (p<0.01) was determined by using a Wilcox rank sum test with Bonferroni correction. Retinal sections were labeled for EdU accumulation (red; **c**) and antibodies to Sox2 (green; **c**). The histogram in **d** illustrates the mean (bar ± SD) and each dot represents one biological replicate. Significance of difference (p-values) was determined by using an unpaired t-test with Bonferroni correction (**d**). Calibration bars in **c** represent 50 µm. Arrows indicate the nuclei of MG.

## Discussion

In this study we investigate how the depletion of microglia from the retina impacts MG and the ability of MG to reprogram into proliferating progenitor-like cells. We found significant transcriptomic changes in resting and activated MG in normal and damaged retinas when the microglia were absent. We identified significant changes in gene modules related to different signaling pathways that have been implicated in regulating the formation of MGPCs. These gene modules included Wnt-, Notch-, fatty acid binding, BMP- and retinoic acid-signaling, which have been implicated in stimulating the proliferation of MGPCs in damaged chick retinas (Campbell et al., 2022; Gallina et al., 2015; Ghai et al., 2010; Hayes et al., 2007; Todd et al., 2017; Todd et al., 2018). Intraocular injections of HBEGF or FGF2 completely rescued the deficit in MGPC formation in damaged retinas missing microglia, whereas injections of RARα agonist or Smad3 antagonist partially rescued this deficit.

It is possible that rapid activation of signaling pathways is required to drive the process of MG reprogramming, and these pathways require rapid activation of signals from microglia. In the mouse retina, for example, microglia upregulate IL1β, IL1α and TNFα within 3 hrs of NMDA-treatment (Todd et al., 2019), and these proinflammatory cytokines activate NFkB signaling in MG (Palazzo et al., 2022; Palazzo et al., 2023). We have recently reported, in the chick retina, that activation of NFkB-signaling with prostratin or injection of a TNF ligand (TNFSF15) rescues the proliferation of MGPCs in damaged retinas where the microglia have been ablated (Palazzo et al., 2020). Prostratin activates NFκB by stimulating phosphorylation and degradation of IκBα in an IKK-dependent manner (Williams et al., 2004). These data indicate that microglia rapidly respond to neuronal damage to produce pro-inflammatory cytokines to stimulate the first steps in MG activation/reactivity that are required for the formation of MGPCs. In fish and chick retinas, MG undergo activation prior to becoming proliferating MGPCs (Hoang et al., 2020). Collectively, these findings suggest that pro-inflammatory cytokines are among the early signals provided by microglia to “kick start” the process of MG reprogramming.

### GSK3β inhibitors fail to stimulate the proliferation of MGPCs

Activation of Wnt-signaling is known to stimulate the proliferation of MGPCs in damaged retinas of zebrafish (Ramachandran et al., 2011), chick (Gallina et al., 2015) and rodents (Osakada et al., 2007; Yao et al., 2016). However, inhibition of GSK3β, which normally occurs downstream of activated Wnt receptors, did not stimulate the proliferation of MGPCs in damaged retinas missing microglia. These results raise the question; why GSK3β inhibition did not stimulate MGPC proliferation in the absence of microglia? It is possible that Wnt-signaling is activated later in the process of reprograming and first requires activation of MAPK signaling initiate the process. Consistent with this notion, HBEGF and components of the MAPK pathway are upregulated within 3 hrs of NMDA-treatment, whereas *WNT4* and *WNT6* are not upregulated until 12hrs after treatment. Alternatively, the targets of GSK3β inhibitors mt be downregulated by MG when microglia are ablated. However, levels of β-catenin and GSK3β are significantly upregulated by MG by in NMDA-damaged retinas, and this upregulation is not affected by the ablation of microglia (not shown). Thus, the targets for GSK3β inhibitors were in place in MG to elicit a response, and this response appears to be restricted to cellular migration.

The delamination of MG nuclei away from the middle of the INL is associated with the proliferation of MGPCs (Fischer and Reh, 2001b; Fischer and Reh, 2003b; Fischer et al., 2004). The vast majority of proliferating MGPCs in M-phase are displaced away from the middle of the INL and are usually found in the proximal INL or distal INL and ONL (Fischer and Reh, 2001b; Fischer and Reh, 2003b). Further, the formation of MGPCs is always associated with delamination of MG nuclei away from the middle of the INL (Fischer and Bongini, 2010; Gallina et al., 2014). We found wide-spread delamination of MG nuclei away from the middle of the INL in response to GSK3β inhibitors in damaged retinas missing microglia, without evidence of proliferation. Similarly, activation of Hedgehog-signaling with a smoothened-agonist and IGF1 stimulates the delamination of MG nuclei without proliferation (Todd and Fischer, 2015). Further, application of MMP2/9 inhibitor with FGF2 stimulates the delamination of MG nuclei (Campbell et al., 2019). By comparison, inhibition of S-adenosyl homocysteine hydrolase and histone methylation potently suppresses the delamination of MG nuclei *and* proliferation in retinas treated with NMDA-damage or insulin+FGF2 (Campbell et al., 2023). Thus, these observations indicate that migration can be de-coupled from proliferation, but proliferation of MGPCs is always associated with nuclear migration.

### Effects of FGF2 and insulin+FGF2

We have previously reported that FGF2 alone or insulin+FGF2 stimulate the formation of proliferating MGPCs in undamaged retinas (Fischer et al., 2002; Fischer et al., 2014). However, these treatments fail to stimulate the formation of MGPCs in undamaged retinas when the microglia have been ablated (Fischer et al., 2014). Interestingly, we found that intraocular injection of FGF2 stimulated the formation of MGPCs in damaged retinas missing microglia. Collectively, these findings suggest that damaged neurons in retinas missing microglia provide signals that permit or act synergistically with FGF2 to drive the reprogramming of MG into proliferating MGPCs. The identity of signals provided by NMDA-damaged retinal neurons remains unknown.

### Amplitude of effects on MGPC rescue

HBEGF and FGF2 had robust rescue effects upon stimulating the proliferation of MGPCs in damaged retinas missing microglia. Consistent with these observations, HBEGF and FGF2 potently stimulate the proliferation of MGPCs in damaged retinas (Fischer et al., 2014; Todd et al., 2015). Similarly, inhibitors of MAPK-signaling and FGF receptors potently suppress the proliferation of MGPCs in damaged retinas (Fischer et al., 2009a; Fischer et al., 2009b). By comparison, Smad3 inhibitor and RAR agonist had relatively small effects upon stimulating the proliferation of MGPCs in damaged retinas missing microglia. Alternatively, it is possible that the failed upregulation of *RARA* by MG in retinas missing microglia was not efficiently rescued because the levels of *RARA* remained low, leaving the agonist with few targets to act upon. Another interpretation is that retinoic acid signaling “fine-tunes” the proliferating stimulating pathways, and the TGFβ/Smad3 acts to suppress the proliferation response.

## Conclusions

We conclude that quiescent and reactive microglia have a significant impact upon the transcriptomic profile of MG. The absence of reactive microglia from damaged retinas results in the failure of activated MG to upregulate many different networks of genes related to different pathways known to influence the reprogramming of MG into proliferating MGPCs. Many of these genes are rapidly up- or downregulated by MG in response to acute retinal damage. We conclude that signals produced by reactive microglia in damaged retinas normally stimulate MG to upregulate cell signaling through HBEGF, FGF/MAPK and RAR, and downregulate inhibitory signaling through TGFβ/Smad3.

## Data availability

Gene-Cell matrices for scRNA-seq data for libraries from saline and NMDA-treated retinas are available through GitHub and OSUMC Sharepoint:

https://github.com/jiewwwang/Singlecell-retinal-regeneration

https://github.com/fischerlab3140/scRNAseq_libraries chick retina scRNA-seq Cell Ranger output files scRNA-Seq data for chick retinas treated with NMDA or insulin and FGF2 can be queried at: https://proteinpaint.stjude.org/F/2019.retina.scRNA.html

## Author Contributions

HE coordinated experiments, performed bioinformatic analyses and contributed to writing the manuscript. AR performed experiments and collected data. WAC and IP generated scRNA-seq libraries and performed bioinformatics. AJF secured funding, designed experiments, analyzed data, constructed figures and wrote the manuscript.

## Acknowledgements

This work was supported by RO1 EY032141-02 and RO1 EY022030-10 (AJF).

---

**Supplemental Figure 1:**
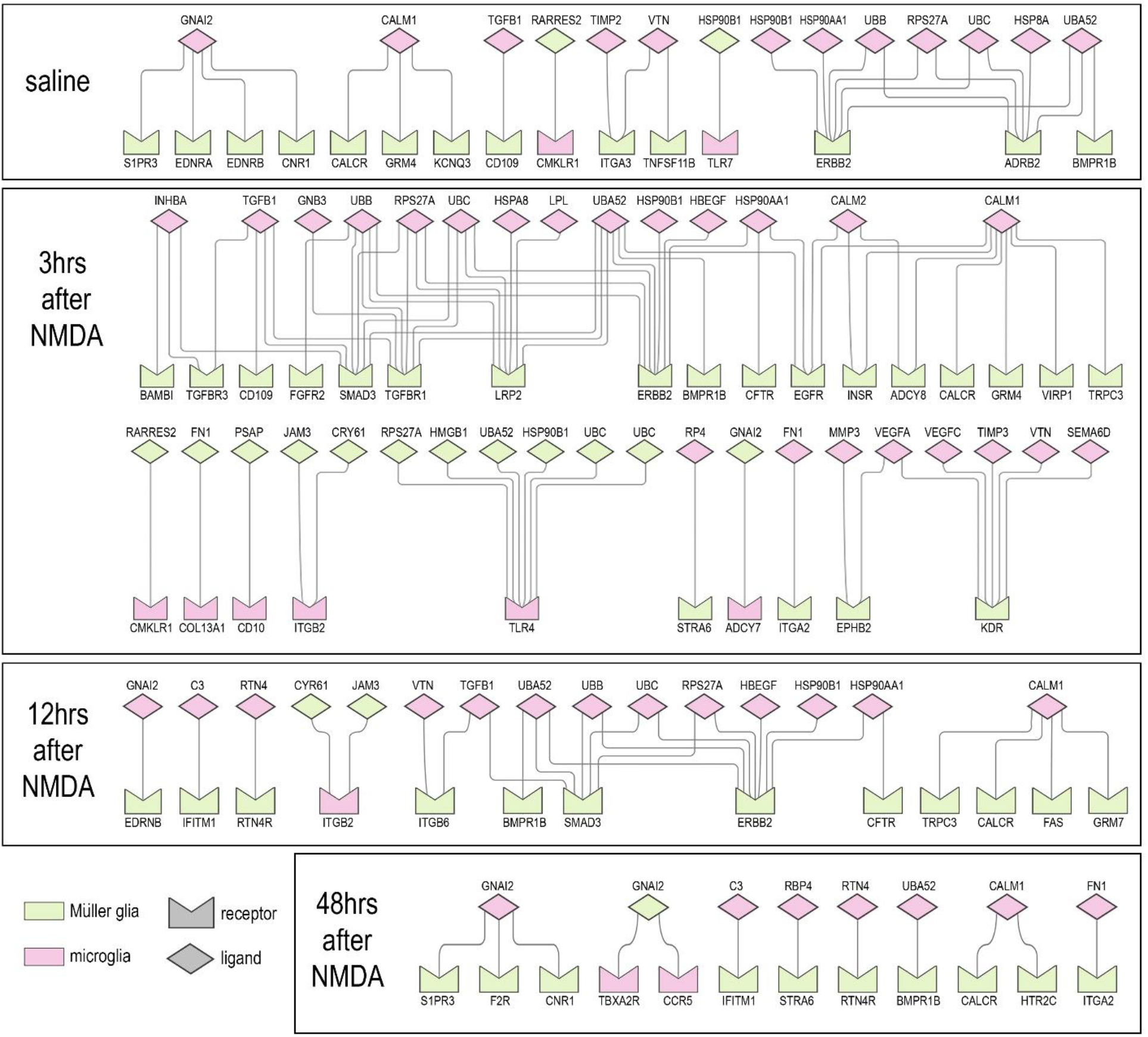
Inferred autocrine ligand-receptor (LR) interactions between MG. scRNA-seq was used to identify putative LR interactions in control and NMDA-damaged retinas at 3, 12 and 48 hours after treatment. SingleCellSignalR was used to identify putative LR interactions. Significant LR-interactions between microglia and MG are illustrated for undamaged and damaged retinas with or without microglia.

## References

Antenos, M., Zhu, J., Jetly, N. M. and Woodruff, T. K. (2008). An activin/furin regulatory loop modulates the processing and secretion of inhibin alpha- and betaB-subunit dimers in pituitary gonadotrope cells. J. Biol. Chem. 283, 33059–33068.

Bernardos, R. L., Barthel, L. K., Meyers, J. R. and Raymond, P. A. (2007). Late-stage neuronal progenitors in the retina are radial Muller glia that function as retinal stem cells. J Neurosci 27, 7028–40.

Butler, A., Hoffman, P., Smibert, P., Papalexi, E. and Satija, R. (2018). Integrating single-cell transcriptomic data across different conditions, technologies, and species. Nat. Biotechnol. 36, 411–420.

Cabello-Aguilar, S., Alame, M., Kon-Sun-Tack, F., Fau, C., Lacroix, M. and Colinge, J. (2020). SingleCellSignalR: inference of intercellular networks from single-cell transcriptomics. Nucleic Acids Res. 48, e55.

Campbell, W. A., Deshmukh, A., Blum, S., Todd, L., Mendonca, N., Weist, J., Zent, J., Hoang, T. V., Blackshaw, S., Leight, J., et al. (2019). Matrix-metalloproteinase expression and gelatinase activity in the avian retina and their influence on Müller glia proliferation. Exp. Neurol. 320, 112984.

Campbell, W. A., Fritsch-Kelleher, A., Palazzo, I., Hoang, T., Blackshaw, S. and Fischer, A. J. (2021a). Midkine is neuroprotective and influences glial reactivity and the formation of Müller glia-derived progenitor cells in chick and mouse retinas. Glia 69, 1515–1539.

Campbell, W. A., Blum, S., Reske, A., Hoang, T., Blackshaw, S. and Fischer, A. J. (2021b). Cannabinoid signaling promotes the de-differentiation and proliferation of Müller glia-derived progenitor cells. Glia 69, 2503–2521.

Campbell, W. A., Tangeman, A., El-Hodiri, H. M., Hawthorn, E. C., Hathoot, M., Blum, S., Hoang, T., Blackshaw, S. and Fischer, A. J. (2022). Fatty acid-binding proteins and fatty acid synthase influence glial reactivity and promote the formation of Müller glia-derived progenitor cells in the chick retina. Dev. Camb. Engl. 149, dev200127.

Campbell, W. A., El-Hodiri, H. M., Torres, D., Hawthorn, E. C., Kelly, L. E., Volkov, L., Akanonu, D. and Fischer, A. J. (2023). Chromatin access regulates the formation of Müller glia-derived progenitor cells in the retina. Glia 71, 1729–1754.

El-Hodiri, H. M., Campbell, W. A., Kelly, L. E., Hawthorn, E. C., Schwartz, M., Jalligampala, A., McCall, M. A., Meyer, K. and Fischer, A. J. (2022). Nuclear Factor I in neurons, glia and during the formation of Müller glia-derived progenitor cells in avian, porcine and primate retinas. J. Comp. Neurol. 530, 1213–1230.

Fausett, B. V. and Goldman, D. (2006). A role for alpha1 tubulin-expressing Muller glia in regeneration of the injured zebrafish retina. J Neurosci 26, 6303–13.

Fausett, B. V., Gumerson, J. D. and Goldman, D. (2008). The proneural basic helix-loop-helix gene ascl1a is required for retina regeneration. J. Neurosci. Off. J. Soc. Neurosci. 28, 1109–1117.

Fischer, A. J. and Bongini, R. (2010). Turning Müller glia into neural progenitors in the retina. Mol. Neurobiol. 42, 199–209.

Fischer, A. J. and Reh, T. A. (2001a). Müller glia are a potential source of neural regeneration in the postnatal chicken retina. Nat. Neurosci. 4, 247–252.

Fischer, A. J. and Reh, T. A. (2001b). Muller glia are a potential source of neural regeneration in the postnatal chicken retina. Nat Neurosci 4, 247–52.

Fischer, A. J. and Reh, T. A. (2003a). Potential of Muller glia to become neurogenic retinal progenitor cells. Glia 43, 70–6.

Fischer, A. J. and Reh, T. A. (2003b). Potential of Muller glia to become neurogenic retinal progenitor cells. Glia 43, 70–6.

Fischer, A. J., Seltner, R. L. P., Poon, J. and Stell, W. K. (1998). Immunocytochemical characterization of quisqualic acid- and N-methyl-D-aspartate-induced excitotoxicity in the retina of chicks. J. Comp. Neurol. 393, 1–15.

Fischer, A. J., Hendrickson, A. and Reh, T. A. (2001). Immunocytochemical characterization of cysts in the peripheral retina and pars plana of the adult primate. Invest Ophthalmol Vis Sci 42, 3256–63.

Fischer, A. J., McGuire, C. R., Dierks, B. D. and Reh, T. A. (2002). Insulin and fibroblast growth factor 2 activate a neurogenic program in Muller glia of the chicken retina. J Neurosci 22, 9387–98.

Fischer, A. J., Schmidt, M., Omar, G. and Reh, T. A. (2004). BMP4 and CNTF are neuroprotective and suppress damage-induced proliferation of Muller glia in the retina. Mol Cell Neurosci 27, 531–42.

Fischer, A. J., Scott, M. A. and Tuten, W. (2009a). Mitogen-activated protein kinase-signaling stimulates Muller glia to proliferate in acutely damaged chicken retina. Glia 57, 166–81.

Fischer, A. J., Scott, M. A., Ritchey, E. R. and Sherwood, P. (2009b). Mitogen-activated protein kinase-signaling regulates the ability of Müller glia to proliferate and protect retinal neurons against excitotoxicity. Glia 57, 1538–1552.

Fischer, A. J., Zelinka, C., Gallina, D., Scott, M. A. and Todd, L. (2014). Reactive microglia and macrophage facilitate the formation of Muller glia-derived retinal progenitors. Glia 62, 1608–28.

Fischer, A. J., Zelinka, C. and Milani-Nejad, N. (2015). Reactive retinal microglia, neuronal survival, and the formation of retinal folds and detachments. Glia 63, 313–27.

Gallina, D., Todd, L. and Fischer, A. J. (2014). A comparative analysis of Müller glia-mediated regeneration in the vertebrate retina. Exp. Eye Res. 123, 121–130.

Gallina, D., Palazzo, I., Steffenson, L., Todd, L. and Fischer, A. J. (2015). Wnt/betacatenin-signaling and the formation of Muller glia-derived progenitors in the chick retina. Dev Neurobiol.

Ghai, K., Zelinka, C. and Fischer, A. J. (2009). Serotonin released from amacrine neurons is scavenged and degraded in bipolar neurons in the retina. J. Neurochem. 111, 1–14.

Ghai, K., Zelinka, C. and Fischer, A. J. (2010). Notch signaling influences neuroprotective and proliferative properties of mature Muller glia. J Neurosci 30, 3101–12.

Guca, E., Suñol, D., Ruiz, L., Konkol, A., Cordero, J., Torner, C., Aragon, E., Martin-Malpartida, P., Riera, A. and Macias, M. J. (2018). TGIF1 homeodomain interacts with Smad MH1 domain and represses TGF-β signaling. Nucleic Acids Res. 46, 9220–9235.

Hayes, S., Nelson, B. R., Buckingham, B. and Reh, T. A. (2007). Notch signaling regulates regeneration in the avian retina. Dev Biol 312, 300–11.

Hitchcock, P. F. and Raymond, P. A. (1992). Retinal regeneration. Trends Neurosci 15, 103– 8.

Hoang, T., Wang, J., Boyd, P., Wang, F., Santiago, C., Jiang, L., Yoo, S., Lahne, M., Todd, L. J., Jia, M., et al. (2020). Gene regulatory networks controlling vertebrate retinal regeneration. Science 370,.

Huang, T., Cui, J., Li, L., Hitchcock, P. F. and Li, Y. (2012). The role of microglia in the neurogenesis of zebrafish retina. Biochem Biophys Res Commun 421, 214–20.

Jorstad, N. L., Wilken, M. S., Grimes, W. N., Wohl, S. G., VandenBosch, L. S., Yoshimatsu, T., Wong, R. O., Rieke, F. and Reh, T. A. (2017). Stimulation of functional neuronal regeneration from Muller glia in adult mice. Nature.

Jouannet, S., Saint-Pol, J., Fernandez, L., Nguyen, V., Charrin, S., Boucheix, C., Brou, C., Milhiet, P.-E. and Rubinstein, E. (2016). TspanC8 tetraspanins differentially regulate the cleavage of ADAM10 substrates, Notch activation and ADAM10 membrane compartmentalization. Cell. Mol. Life Sci. CMLS 73, 1895–1915.

Karl, M. O., Hayes, S., Nelson, B. R., Tan, K., Buckingham, B. and Reh, T. A. (2008). Stimulation of neural regeneration in the mouse retina. Proc Natl Acad Sci U A 105, 19508–13.

LeBaron, R. G., Bezverkov, K. I., Zimber, M. P., Pavelec, R., Skonier, J. and Purchio, A. F. (1995). Beta IG-H3, a novel secretory protein inducible by transforming growth factor-beta, is present in normal skin and promotes the adhesion and spreading of dermal fibroblasts in vitro. J. Invest. Dermatol. 104, 844–849.

Lemjabbar, H. and Basbaum, C. (2002). Platelet-activating factor receptor and ADAM10 mediate responses to Staphylococcus aureus in epithelial cells. Nat. Med. 8, 41–46.

Ling, N., Ying, S. Y., Ueno, N., Shimasaki, S., Esch, F., Hotta, M. and Guillemin, R. (1986). Pituitary FSH is released by a heterodimer of the beta-subunits from the two forms of inhibin. Nature 321, 779–782.

Mitra, S., Devi, S., Lee, M.-S., Jui, J., Sahu, A. and Goldman, D. (2022). Vegf signaling between Müller glia and vascular endothelial cells is regulated by immune cells and stimulates retina regeneration. Proc. Natl. Acad. Sci. U. S. A. 119, e2211690119.

Ooto, S., Akagi, T., Kageyama, R., Akita, J., Mandai, M., Honda, Y. and Takahashi, M. (2004). Potential for neural regeneration after neurotoxic injury in the adult mammalian retina. Proc Natl Acad Sci U A 101, 13654–9.

Osakada, F., Ooto, S., Akagi, T., Mandai, M., Akaike, A. and Takahashi, M. (2007). Wnt signaling promotes regeneration in the retina of adult mammals. J Neurosci 27, 4210–9.

Palazzo, I., Deistler, K., Hoang, T. V., Blackshaw, S. and Fischer, A. J. (2020). NF-κB signaling regulates the formation of proliferating Müller glia-derived progenitor cells in the avian retina. Dev. Camb. Engl. 147, dev183418.

Palazzo, I., Todd, L. J., Hoang, T. V., Reh, T. A., Blackshaw, S. and Fischer, A. J. (2022). NFkB-signaling promotes glial reactivity and suppresses Müller glia-mediated neuron regeneration in the mammalian retina. Glia 70, 1380–1401.

Palazzo, I., Kelly, L., Koenig, L. and Fischer, A. J. (2023). Patterns of NFkB activation resulting from damage, reactive microglia, cytokines, and growth factors in the mouse retina. Exp. Neurol. 359, 114233.

Pollak, J., Wilken, M. S., Ueki, Y., Cox, K. E., Sullivan, J. M., Taylor, R. J., Levine, E. M. and Reh, T. A. (2013). Ascl1 reprograms mouse Muller glia into neurogenic retinal progenitors. Development 140, 2619–2631.

Ramachandran, R., Zhao, X. F. and Goldman, D. (2011). Ascl1a/Dkk/beta-catenin signaling pathway is necessary and glycogen synthase kinase-3beta inhibition is sufficient for zebrafish retina regeneration. Proc Natl Acad Sci U A 108, 15858–63.

Raymond, P. A. (1991). Retinal regeneration in teleost fish. Ciba Found Symp 160, 171–86; discussion 186-91.

Reichenbach, A. and Bringmann, A. (2013). New functions of Müller cells. Glia 61, 651–678.

Ritchey, E. R., Bongini, R. E., Code, K. A., Zelinka, C., Petersen-Jones, S. and Fischer, A. J. (2010). The pattern of expression of guanine nucleotide-binding protein beta3 in the retina is conserved across vertebrate species. Neuroscience 169, 1376–91.

Satija, R., Farrell, J. A., Gennert, D., Schier, A. F. and Regev, A. (2015). Spatial reconstruction of single-cell gene expression data. Nat Biotechnol 33, 495–502.

Seegar, T. C. M., Killingsworth, L. B., Saha, N., Meyer, P. A., Patra, D., Zimmerman, B., Janes, P. W., Rubinstein, E., Nikolov, D. B., Skiniotis, G., et al. (2017). Structural Basis for Regulated Proteolysis by the α-Secretase ADAM10. Cell 171, 1638–1648.e7.

Shannon, P., Markiel, A., Ozier, O., Baliga, N. S., Wang, J. T., Ramage, D., Amin, N., Schwikowski, B. and Ideker, T. (2003). Cytoscape: a software environment for integrated models of biomolecular interaction networks. Genome Res. 13, 2498–2504.

Skonier, J., Bennett, K., Rothwell, V., Kosowski, S., Plowman, G., Wallace, P., Edelhoff, S., Disteche, C., Neubauer, M. and Marquardt, H. (1994). beta ig-h3: a transforming growth factor-beta-responsive gene encoding a secreted protein that inhibits cell attachment in vitro and suppresses the growth of CHO cells in nude mice. DNA Cell Biol. 13, 571–584.

Todd, L. and Fischer, A. J. (2015). Hedgehog-signaling stimulates the formation of proliferating Müller glia-derived progenitor cells in the retina. Development 142, 2610– 2622.

Todd, L., Volkov, L. I., Zelinka, C., Squires, N. and Fischer, A. J. (2015). Heparin-binding EGF-like growth factor (HB-EGF) stimulates the proliferation of Muller glia-derived progenitor cells in avian and murine retinas. Mol Cell Neurosci 69, 54–64.

Todd, L., Palazzo, I., Squires, N., Mendonca, N. and Fischer, A. J. (2017). BMP- and TGFbeta-signaling regulate the formation of Muller glia-derived progenitor cells in the avian retina. Glia.

Todd, L., Suarez, L., Quinn, C. and Fischer, A. J. (2018). Retinoic Acid-Signaling Regulates the Proliferative and Neurogenic Capacity of Muller Glia-Derived Progenitor Cells in the Avian Retina. Stem Cells 36,.

Todd, L., Palazzo, I., Suarez, L., Liu, X., Volkov, L., Hoang, T. V., Campbell, W. A., Blackshaw, S., Quan, N. and Fischer, A. J. (2019). Reactive microglia and IL1β/IL-1R1-signaling mediate neuroprotection in excitotoxin-damaged mouse retina. J. Neuroinflammation 16, 118.

Todd, L., Finkbeiner, C., Wong, C. K., Hooper, M. J. and Reh, T. A. (2020). Microglia Suppress Ascl1-Induced Retinal Regeneration in Mice. Cell Rep. 33, 108507.

Ueki, Y., Wilken, M. S., Cox, K. E., Chipman, L., Jorstad, N., Sternhagen, K., Simic, M., Ullom, K., Nakafuku, M. and Reh, T. A. (2015). Transgenic expression of the proneural transcription factor Ascl1 in Muller glia stimulates retinal regeneration in young mice. Proc Natl Acad Sci U A 112, 13717–22.

Wan, J. and Goldman, D. (2016). Retina regeneration in zebrafish. Curr Opin Genet Dev 40, 41–47.

Wan, J., Ramachandran, R. and Goldman, D. (2012). HB-EGF is necessary and sufficient for Muller glia dedifferentiation and retina regeneration. Dev Cell 22, 334–47.

White, D. T., Sengupta, S., Saxena, M. T., Xu, Q., Hanes, J., Ding, D., Ji, H. and Mumm, J. S. (2017). Immunomodulation-accelerated neuronal regeneration following selective rod photoreceptor cell ablation in the zebrafish retina. Proc. Natl. Acad. Sci. U. S. A. 114, E3719–E3728.

Williams, S. A., Chen, L.-F., Kwon, H., Fenard, D., Bisgrove, D., Verdin, E. and Greene, W. C. (2004). Prostratin antagonizes HIV latency by activating NF-kappaB. J. Biol. Chem. 279, 42008–42017.

Yao, K., Qiu, S., Tian, L., Snider, W. D., Flannery, J. G., Schaffer, D. V. and Chen, B. (2016). Wnt Regulates Proliferation and Neurogenic Potential of Muller Glial Cells via a Lin28/let-7 miRNA-Dependent Pathway in Adult Mammalian Retinas. Cell Rep 17, 165– 78.

Zelinka, C. P., Scott, M. A., Volkov, L. and Fischer, A. J. (2012). The Reactivity, Distribution and Abundance of Non-Astrocytic Inner Retinal Glial (NIRG) Cells Are Regulated by Microglia, Acute Damage, and IGF1. PLoS One 7, e44477.

